# Mobile Eye-Tracking Glasses Capture Ocular and Head Markers of Listening Effort

**DOI:** 10.1101/2025.09.17.676957

**Authors:** M. Eric Cui, Emilie Verno-Lavigne, Shreshth Saxena, Lauren K. Fink, Björn Herrmann

## Abstract

To extend the assessment of listening effort beyond a sound booth, we validated mobile eye-tracking glasses (Pupil Labs Neon) by comparing them to a stationary system (Eyelink DUO) in a controlled environment. We recorded eye movements, pupil size, and head movements from 26 young adults during a speech-in-noise task. When listening conditions became challenging, we observed reduced gaze dispersion and increased pupil sizes of similar magnitude from both devices, in addition to reduced head movements recorded solely by the mobile device. These findings suggest that mobile eye-trackers reliably capture listening effort, paving the path towards assessments in daily settings.

## 1. Introduction

Comprehending speech in noisy environments, such as a crowded restaurant, places high cognitive demand on attentional and memory resources, requiring listening effort (Herrmann & Johnsrude, 2020; Peelle, 2018; Pichora-Fuller et al., 2016a). Such effort is often considered an early indicator of hearing loss, making its objective assessment critical for clinical and research purposes (Helfer & Jesse, 2021; Herrmann & Johnsrude, 2020; Pichora-Fuller et al., 2016). To date, pupillometry is one of the most commonly used objective markers to listening effort, with numerous studies demonstrating that pupil size increases as listening becomes more demanding (Almaraz et al., 2018; Piquado et al., 2010; Van Der Wel & Van Steenbergen, 2018; Winn et al., 2018a; Zekveld et al., 2018). More recently, a separate line of work indicates that a reduction in eye movements may also index listening effort, potentially through pupil-independent mechanisms (Cui & Herrmann, 2023; Herrmann & Ryan, 2024). Assessing listening effort using eye movements offers a distinct advantage compared to pupillometry: eye-based metrics are less sensitive to environmental factors like ambient lighting, stimulus luminance, and baseline physiological fluctuations (Fink et al., 2024; Thurman et al., 2021; Winn et al., 2018a; Zekveld et al., 2018). This makes eye movements a promising candidate for assessing listening effort in more natural, real-world situations. However, both pupillometry and eye-movement research have mainly relied on stationary eye-trackers that require restricted head movements, limiting the ecological validity of these assessments.

Modern mobile eye-tracking glasses present an opportunity to overcome these limitations and move assessments towards real-world, out-of-lab contexts. These portable devices have been successfully used to track eye movements and pupil size across different age groups and hearing abilities (Chavant & Kapoula, 2022; Hidalgo et al., 2023; Perea Perez, 2023) for various auditory tasks, such as music listening and performing (Saxena et al., 2025; Vandemoortele et al., 2018), sound localization (Alemu et al., 2024), and speech-in-noise perception (Perea Perez, 2023). This evidence suggests that mobile eye-trackers could effectively measure listening effort. For example, promising correspondence between mobile eye-tracking glasses and a stationary EyeLink eye-tracker (the most commonly used system to achieve high temporal and spatial precision) has been found for simple oculomotor tasks (Ehinger et al., 2019). However, it remains unclear whether relative measures, such as gaze dispersion (Cui & Herrmann, 2023; Herrmann et al., 2025; Herrmann & Ryan, 2024), a sensitive metric capturing changes in eye movements is robust enough to concurrent reduction in both spatial and temporal resolution.

Beyond expanding previously established listening effort measures from stationary to mobile devices, mobile eye-trackers provide additional unique metrics that may be relevant to listening effort. For instance, head movements (e.g., measured via inertial motion units embedded in head-mounted eye-trackers) may index cognitive load in a manner like eye movements, however, this has not been examined. Listeners reduce eye movements during difficult listening, possibly to limit distracting visual input (Cui & Herrmann, 2023). A similar pattern might occur for head movements, where a reduction in head movements might reduce distracting proprioceptive inputs, in turn freeing up cognitive resources for a listening task (Higgins et al., 2023). This would also be in line with research showing coupling between sensorimotor and cognitive systems (Dault et al., 2001; Craighero 2022). Integrating these markers could provide a comprehensive and ecologically valid approach to monitoring the dynamics of listening effort through mobile glasses.

The present study examines whether modern mobile eye-tracking glasses (Pupil Labs’ Neon) are similarly sensitive to pupil-size and eye-movement markers of listening effort as a stationary eye-tracking system (EyeLink Duo). We also investigate whether head movements index listening effort. To this end, we concurrently measured pupil size, gaze dispersion, and head-movement variability with synchronized stationary and mobile devices concurrently while younger, normal-hearing adults performed a speech-comprehension task under easy and difficult speech-masking conditions.

## 2. Methods

### 2.1 Participants

Twenty-six younger adults (M = 25.75 years, SD = 5.25, range = 18-35 years; 17 females, 6 males, 1 agender, 2 non-binary) participated in the study. All self-reported being native English speakers, having normal vision and hearing, and having no severe psychological or neurological conditions. All participants provided written informed consent and received $20 per hour for their participation. Two additional individuals participated, but their data were not recorded (N = 1) or the data timestamp procedure failed (N = 1). The study was approved by the Research Ethics Board at Baycrest Academy for Research and Education.

### 2.2 Experiment Setup and Procedure

Participants sat in a sound-attenuating booth with their head stabilized on a chin and forehead rest, positioned approximately 70 cm from a computer monitor. Auditory stimuli were presented binaurally through Sony MDR-7506 headphones, with sound played through a Steinberg UR22 mkII external sound card. The experiment was programmed in MATLAB using Psychtoolbox (v3.0.14). The main task was a speech-in-noise perception task. Participants listened to 180 Harvard sentences (mean duration = 2.5 s; Institute of Electrical and Electronics Engineers, 1969) presented in a 7-s 12-talker babble noise (Bilger, 1984), either under an easy (+11 dB SNR) or difficult (-4 dB SNR) listening condition. The sentence started 1.5-s after babble onset and the SNR was determined by adjusting the level of the speech, whereas the level of the babble was constant across trials (Kadem et al., 2020; Ohlenforst et al., 2017). The computer screen remained blank before and during sound presentation. After each sentence, a probe word appeared on the computer screen, and participants judged whether it was semantically related to the preceding sentence (Cui & Herrmann, 2023; Kadem et al., 2020). Participants completed 5 blocks (although 3 participants missed 1, 2, or 3 blocks of data due to incidental technical issues). Each block comprised 36 trials (18 per SNR condition), and trials were pseudo-randomized such that a maximum of 2 trials of the same SNR could occur in direct succession. A 10-trial practice block preceded the experiment. Behavioral performance in the semantic-relatedness task was calculated as the proportion of correct responses. A paired-samples t-test was used to assess the difference between easy and difficult SNRs.

### 2.3 Recording of Stationary and Mobile Eye-Trackers

To enable a direct comparison of the two eye-tracking systems, we recorded participants’ eye movements and pupil size concurrently using both a stationary and a mobile eye-tracker. An Eyelink DUO was used as a high-precision stationary eye-tracker to record eye movements and pupil size for both eyes at a sampling rate of 1000 Hz. We refer to this device as the “stationary” eye tracker hereafter. A standard nine-point calibration and validation procedure (McIntire et al., 2014) was performed at the beginning of each experimental block. A timestamp message was sent to the eye tracker at the beginning of each babble onset. We used Pupil Labs Neon mobile eye-tracking glasses with a Motorola Edge 40 Pro Android smartphone running the Neon Companion app (v2.8.2) (Baumann & Dierkes, 2023) for data collection and management. We refer to this device as the “mobile” eye tracker hereafter. Eye-movement and pupil-size data were recorded at 200 Hz, whereas head movement data were collected at 110 Hz. The calibration for the mobile glasses involved two steps. First, each participant’s intraocular distance was measured and entered into the recording app. Second, participants completed a brief, single-point calibration by fixating on a designated target. On the laptop that was used for stimulus presentation, we also used SocialEyes software written in Python (Saxena et al., 2025) to remotely control the smart phone and Neon Companion app (e.g., start and stop recording). We used SocialEyes’ timestamp synchronization procedure to estimate the offset between the internal clocks of the stimulus computer and the recording smartphone (Saxena et al., 2025). Offsets between the two devices clocks were estimated approximately every 10-s during data collection for accurate timestamp calculation during offline analysis.

### 2.4 Eye-movement, pupil-size, and head-movement data processing

Data were processed offline using MATLAB and Python scripts. The x-coordinate, y-coordinate, and pupil-size data from the stationary eye-tracker were down-sampled to 200 Hz to match the sampling frequency of the eye data from the mobile eye-tracker. Mobile eye-tracking data were corrected for lens distortion using built-in functions from the OpenCV library (Bradski, 2000). Timestamps were estimated through linear interpolation using the recorded offsets between the independent internal clocks of the stimulus computer and the recording smartphone (Saxena et al., 2025). The remaining steps for data processing were similar for both eye trackers.

Pupil-size and x- and y-coordinate data were set to *NaN* (not a number in MATLAB) for 100-ms before and 200-ms after each blink, as well as any outliers that deviated by more than 3 standard deviations from the participant’s block mean. All removed and missing data points (*NaN*-coded) were interpolated using a shape-preserving piecewise cubic interpolation (*pchip*). Pupil-size data was then low-pass filtered at 5 Hz (FIR, 131 points, Kaiser Window).

We calculated gaze dispersion as a broad measure of eye movements. We recently developed this measure to non-specifically capture any eye-movement changes associated with listening effort (Cui & Herrmann, 2023; Herrmann et al., 2025; Herrmann & Ryan, 2024). Gaze dispersion was calculated as the log-transformed standard deviation of gaze coordinates (averaged across x and y) within a 1-s sliding window (ignoring data points that had been NaN-coded during preprocessing; and requiring at least 20 data points within a 1-s time window). A smaller value indicates fewer eye movements.

The continuous pupil-size and gaze-dispersion data were divided into epochs ranging from -1 s to 7 s time-locked to babble onset. The percentage of data points within the -1 to 7 epoch that were marked as NaN during pre-processing did not differ between the stationary device (*M* = 21.49%, *SD* = 19.24%) and the mobile device (M = 21.64%, SD = 13.57%), *t*(25) = 0.029, *p* = 0.977. Epochs containing equal to or over 50% of samples that were marked as NaN during pre-processing were excluded from further analysis (stationary: *M* = 13.27, SD = 18.57%; mobile: M = 9.58%, SD = 16.29%; *t*(25) = 0.677, *p* = 0. 505).

Epochs were averaged separately for easy and difficult SNRs. To enable cross-device comparisons (stationary vs mobile), both pupil-size and gaze-dispersion time courses (using across-trial average) were converted to within-participant z-scores, computed separately for the stationary and mobile devices to account for scaling differences between devices. The pupil-size time courses were then baseline-corrected by subtracting the mean in the pre-babble period [-1, 0] from all data points, separately for each SNR condition (Mathôt et al., 2018; Winn et al., 2018b). Gaze-dispersion data do not require baseline correction. Gyroscope data from the mobile glasses were also analyzed to assess SNR effects on head movements (gyroscope data are not available from stationary eye trackers). To this end, head-movement data were analyzed similarly to gaze dispersion as positional variability, calculating the log-transformed standard deviation of the gyroscope values (yaw, pitch, and roll) within a 1-s sliding time window. A lower value indicates fewer head movements. Positional-variability data were divided into epochs ranging from -1 s to 7 s time-locked to babble onset, and averaged across trials for each SNR condition.

For statistical analyses, gaze dispersion and positional variability were averaged within the 1.5 s to 4 s (1.5 s corresponding to sentence onset; 4 s corresponding to the average sentence offset), separately for the easy and difficult listening conditions. For the pupil size, data were averaged within the 2 s to 4.5 s to account for the known delay in the pupil response (Knapen et al., 2016; Winn et al., 2018a, 2018b; Zhang et al., 2022). Both pupil dilation and gaze dispersion were analyzed using a repeated-measures analysis of variance (rmANOVA). This analysis included two factors: SNR (listening condition: easy vs. difficult) and Eye Tracker (stationary vs. mobile). Pearson correlations were used to evaluate the consistency between the two data from the two eye-tracking devices. A paired-sample t-test was used to assess SNR effect on positional variability (head movements). Effect sizes for ANOVA are reported as omega squared (*ω* ^*2*^), and for t-tests, as Cohen’s d (*d*). All analyses were conducted using MATLAB and JASP software (version 0.95.1). Note that data from only 24 participants were available for the analysis of the pupil size. Pupil-size data from two participants were not successfully uploaded to the Pupil Cloud (https://cloud.pupil-labs.com; where initial Pupil Labs processing is calculated), and could also not be recovered offline.

## 3. Results

### 3.1 Behavioral Performance

As expected, behavioral performance accuracy was lower for the difficult (*M* = 0.77, *SD* = 0.08) than the easy SNR condition (*M* = 0.93, *SD* = 0.05; *t*(25) = 14.30, *p* = 2.78 · 10^-13^, *d* = 2.8).

### 3.2 Pupil Size

The pupil size was larger in the difficult than in the easy condition (SNR effect: *F*(1, 23) = 10.12, *p* = 0.004, *ω*^2^ = 0.12; Figure 1B). The main effect of Eye Tracker was not significant, *F*(1, 23) = 3.66, *p* = 0.068, nor was the interaction, *F*(1, 23) = 1.46, *p* = 0.238, suggesting that the SNR effect on pupil size did not differ between eye-tracking devices (see also Figure 1C). We confirmed that the SNR effect was indeed significant for both devices (stationary: *t*(23) = 4.33, *p* = 2.5· 10^-4^, *d* = 0.88; mobile: *t*(23) = 2.285, *p* = 0.032, *d* = 0.47). Pearson correlations indicated strong associations between the two eye trackers in both the easy (*r* = 0.74, *p* < 0.001) and difficult (*r* = 0.88, *p* < 0.001) listening conditions, suggesting high consistency in pupil size measurements across devices (see Figure 1D).

**Fig. 1.**
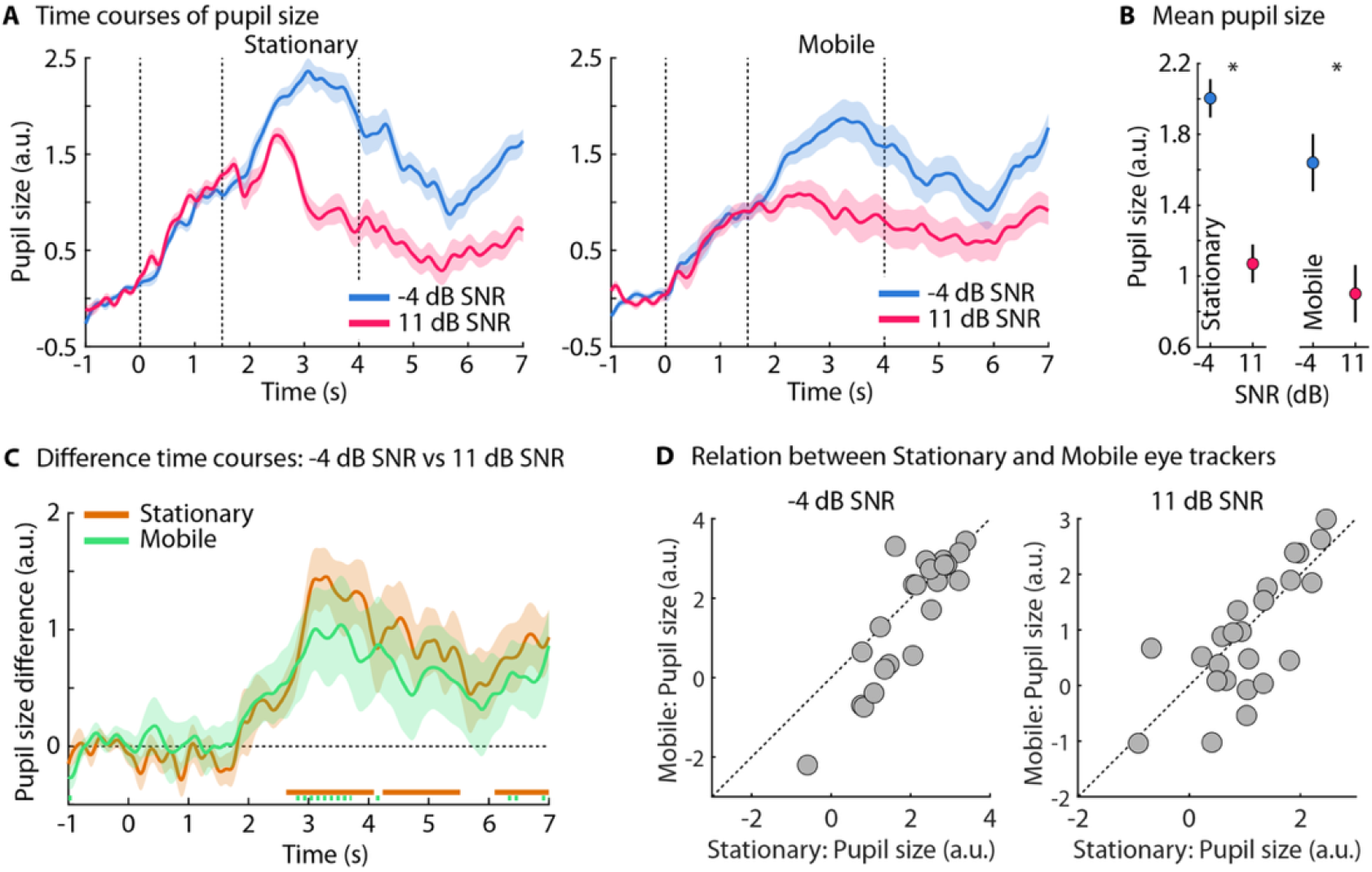
Pupil responses to listening conditions for stationary and mobile eye trackers. (A) Time courses of pupil size (in z□score (within participant); arbitrary units; a.u.) for stationary (left) and mobile (right) eye trackers. Data are presented for difficult (-4 dB SNR; blue) and easy (+11 dB SNR; pink) listening conditions, with the corresponding shaded areas representing standard error of the mean (between-participant variance removed; (Masson et al., 2003)). Dashed lines denote babble onset (0 s), the sentence onset (1.5 s) and average sentence offset (4 s). (B) Mean pupil size (averaged across 2-4.5 s) for each condition and device. Asterisks denote statistical significance (*p* < .05). (C) Time course of the difference in pupil size between SNR conditions for the stationary (orange) and mobile (green) eye trackers. Horizontal lines at the bottom indicate time points of a statistically significant difference from zero (i.e., significant SNR effect; *p* < .05; solid line = corrected using False Discovery Rate (Benjamini & Hochberg, 1995); dashed line = uncorrected). (D) Scatter plots show the correlation between stationary and mobile eye trackers for pupil size during difficult (left) and easy (right) conditions. Data points above the diagonal line indicate that the mobile eye tracker measured a larger pupil size, while points below the line show a larger pupil size measured by the stationary eye tracker.

### 3.3 Gaze Dispersion

Gaze dispersion was lower for the difficult compared to the easy listening condition (SNR effect: *F*(1,25) = 118.77, *p* = 2.92 · 10^-11^, *ω*^*2*^ = 0.52; Figure 2). The effect of Eye Tracker was not significant (*F*(1,25) = 0.28, *p* = 0.605), nor the interaction (*F*(1,25) = 1.30, *p* = 0.265), suggesting gaze dispersion is similarly sensitive to SNR for both device types. We confirmed that the gaze-dispersion effect was significant for the stationary (*t*(25) = 9.89, *p* = 1.47 · 10^-10^, *d* = 1.94) and the mobile eye tracker (*t*(25) = 7.74, *p* = 1.01 · 10^-7^, *d* = 1.52; Figure 2C). Pearson correlations indicated moderate-to-strong correlations between the two eye trackers for both the easy (*r* = 0.58, *p* = 0.003) and difficult SNR (*r* = 0.73, *p* = 2.6 · 10^-6^; Figure 2D).

**Fig. 2.**
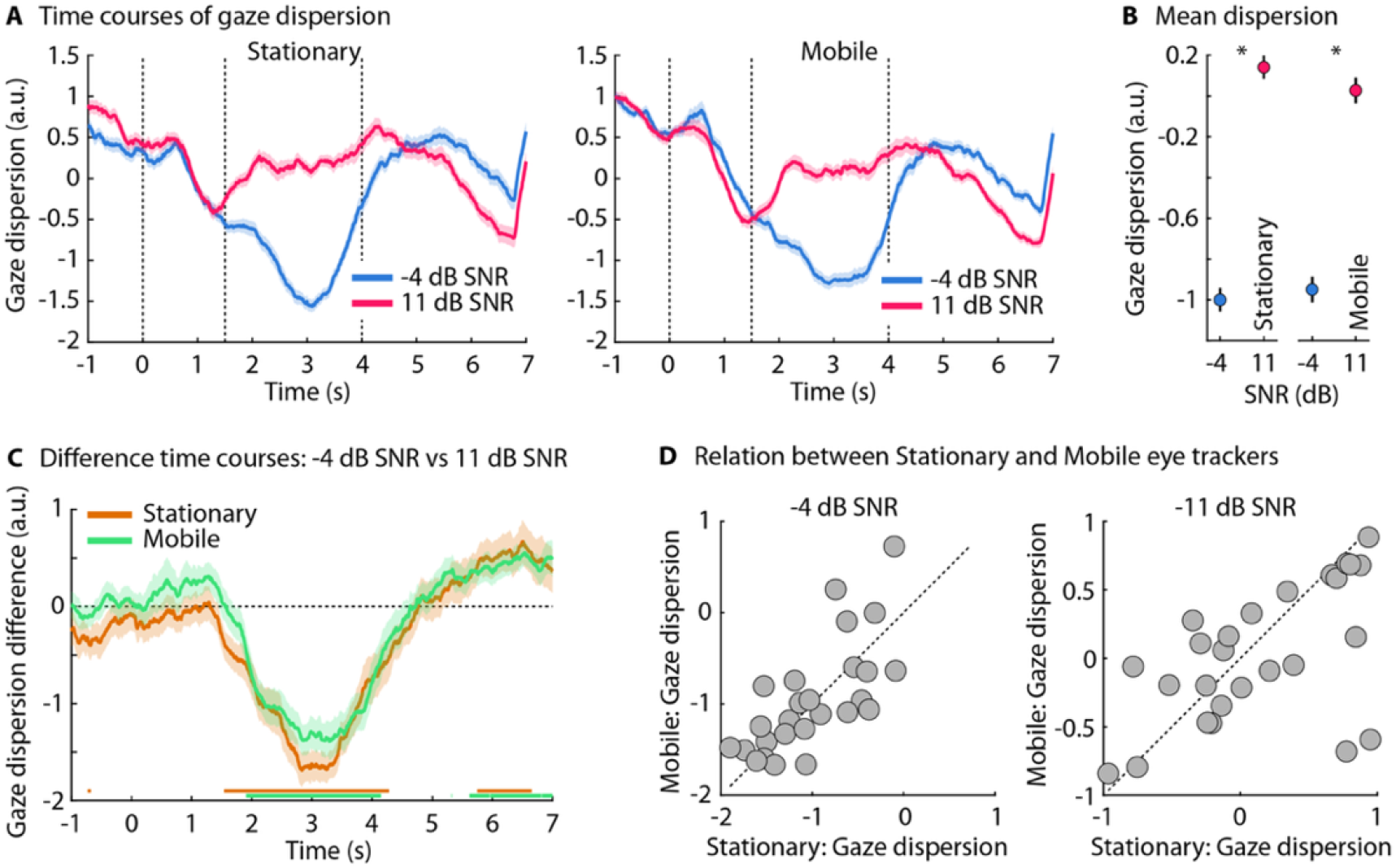
Gaze dispersion responses to listening conditions for stationary and mobile eye trackers. (A) Time courses of gaze dispersion (in z□score (within participant); arbitrary units; a.u.) for stationary (left) and mobile (right) eye trackers. Data are presented for difficult (-4 dB SNR; blue) and easy (+11 dB SNR; pink) listening conditions, with the corresponding shaded areas representing standard error of the mean (between-participant variance removed; (Masson et al., 2003)). Dashed lines denote babble onset (0 s), the sentence onset (1.5 s) and average sentence offset (4 s). (B) Mean gaze dispersion (averaged across 1.5-4 s) for each condition and device. Asterisks denote statistical significance (*p* < .05). (C) Time course of the difference in gaze dispersion between two conditions for the stationary (orange) and mobile (green) eye trackers. Horizontal lines at the bottom indicate time points of a statistically significant difference from zero (i.e., a significant SNR effect; *p* < .05 corrected using False Discovery Rate (Benjamini & Hochberg, 1995)). (D) Scatter plots show the correlation between stationary and mobile eye trackers for gaze dispersion during difficult (left) and easy (right) conditions. Data points above the diagonal line indicate that the mobile eye tracker measured a larger gaze dispersion, while points below the line show a larger gaze dispersion measured by the stationary eye tracker.

### 3.4 Head Movements

Head movements – that is, positional variability – measured using the mobile eye-tracking glasses was lower for the difficult compared to the easy SNR (*t*(25) = 2.97, *p* = 0.006, *d* = 0.58; Figure 3).

**Figure 3.**
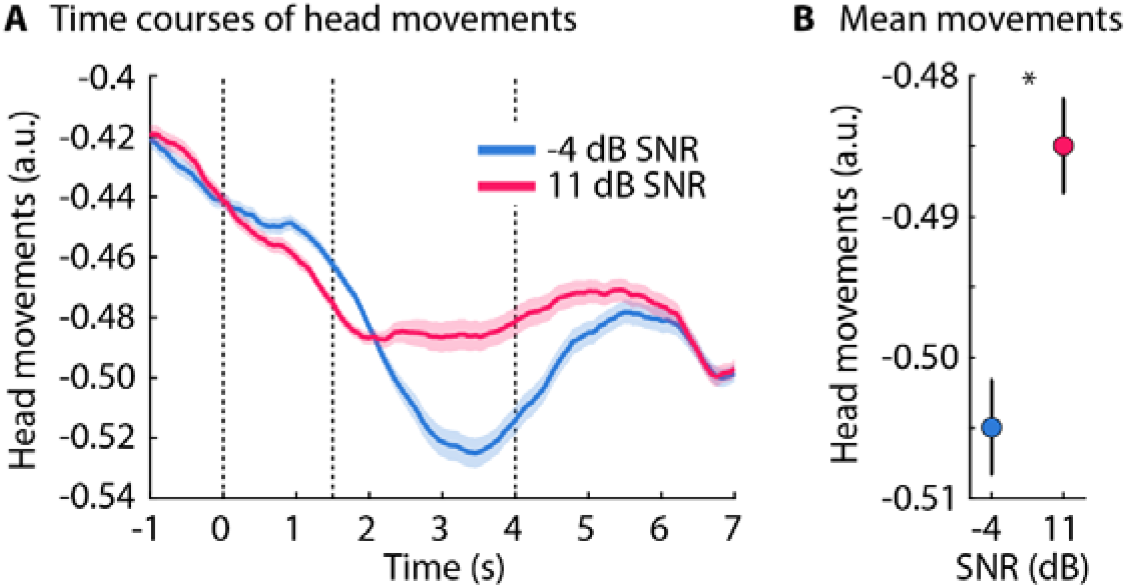
Positional variability of head movements, measured by mobile eye-tracking glasses, index of listening effort. (A) Time courses of positional variability of head movements (in arbitrary units; a.u.). Data are presented for the difficult (-4 dB SNR; blue) and easy (+11 dB SNR; pink) listening conditions. Shaded regions represent the standard error of the mean (between-participant variance removed; (Masson et al., 2003)). Vertical dashed lines mark the babble onset (0 s), sentence onset (1.5 s) and average sentence offset (4 s). (B) Mean head movements (averaged across 1.5-4 s) for each condition. Lower values on the y-axis indicate more stable head position. \**p* < .05.

### 3.5 Relationships Between Listening Effort Measures

Correlation analyses were conducted using the difference scores between the difficult and easy conditions for each measure (i.e., SNR effect). There were no significant relationships between the listening effort markers: SNR effects for gaze dispersion and pupil dilation were not correlated for stationary (*r* = -0.20, *p* = 0.340, 95 % *CI* = [-0.56, 0.22]) nor mobile eye-tracking data (*r* = 0.04, *p* = 0.860, 95 % *CI* = [-0.37, 0.44]). Further analyses using the data from the mobile eye-tracking glasses revealed a marginally significant correlation between positional variability (head movement) and gaze dispersion (*r* = 0.39, *p* = 0.051, 95 % *CI* = [-0.016, 0.69]) and a significant negative correlation with pupil dilation (r = -0.44, *p* = 0.026, 95 % *CI* = [0.045, 0.72]). Noteworthy, these correlation analyses were post-hoc, therefore, they might be underpowered due to a small sample size (N = 24).

## 4. Discussion

The current study investigated whether a pair of state-of-the-art mobile eye-tracking glasses are sensitive to several physiological markers of listening effort. We found reduced gaze dispersion, increased pupil size, and reduced head movements when speech masking – and thus listening effort – increased. Cross-device comparisons found no significant differences in the speech-masking effect, although there was some indication that the pupil-size effect may be numerically smaller for the mobile than the stationary eye-tracker. Overall, the current results indicate high feasibility of using mobile eye-tracking glasses for the assessment of listening effort, including head-movement measures that are not available using a stationary eye-tracker.

### 4.1 Ocular and Head Markers

Both eye-trackers converged on the same pattern: as listening demands increased, gaze dispersion decreased, whereas pupil size increased. These results are highly consistent with previous studies using pupillometry (Almaraz et al., 2018; Piquado et al., 2010; Van Der Wel & Van Steenbergen, 2018; Zekveld et al., 2018) and eye movements (Contadini-Wright et al., 2023; Cui & Herrmann, 2023; Herrmann et al., 2025; Herrmann & Ryan, 2024) to assess listening effort. An increase in pupil size and a reduction in eye movements appear to capture complementary aspects of listening effort (see also (Liu & Chait, 2025)). These measures likely tap into interconnected functions within overlapping networks. For example, pupil size is linked to arousal involving the locus coeruleus-norepinephrine systems (Joshi et al., 2016), which is subject to top-down regulation from prefrontal control regions (Joshi & Gold, 2020; Ross & Van Bockstaele, 2021). Meanwhile, gaze dispersion may index effort through attentional control that suppresses visual exploration, possibly relying on involvement of superior colliculus, thalamus, basal ganglia, cerebellum, and the prefrontal cortex (Lencer et al., 2019; Pierce et al., 2019; Schneider et al., 2014; Sparks, 2002).

Despite the use of a chin and forehead rest, participants still showed reduced head movements as the listening task became more challenging. These results show that head movements persist despite the use of a chin and forehead rest, and it still could be modulated by task demands. Furthermore, the reduction in head movements during effortful listening suggests that individuals may have adopted a freezing or stabilization strategy to free up cognitive resources. By reducing head movements, this strategy could minimize the amount of proprioceptive information (Dault et al., 2001), thereby freeing up more cognitive resources for processing acoustic information in the speech-in-noise task (Assländer & Peterka, 2014). This principle is further supported by neurophysiological evidence from rodents, which shows that reduced body movement is linked to heightened neuronal activity in the auditory cortex (Schneider et al., 2014). Critically, although we observed a marginal correlation between head movements and eye-tracking measures (pupil-size and gaze dispersion), it is unlikely that the SNR effects on gaze dispersion and pupil size are simple artifacts of head movements. For example, the mobile eye-tracking glasses are head-mounted, and the recorded eye movements should thus be unaffected by head movements. Nevertheless, the relationship between head movement and ocular changes presents an interesting area for future research.

### 4.2 Device Comparison

The effect of SNR (speech masking) on gaze dispersion was comparable across the stationary and the mobile eye tracker. For pupil size, the SNR effect was numerically smaller for the mobile compared to the stationary eye-tracker. However, neither the Device main effect nor the Device × SNR interaction was significant, thus providing no formal evidence for a difference between devices. Without over-interpreting the numerically smaller SNR effect for the mobile glasses, potential contributors could be the mobile system’s less precise calibration, stationary geometry vs head-worn geometry for estimating the pupil size (Brisson et al., 2013; Hayes & Petrov, 2016), or generally a lower sensitivity. Calibration for the mobile glasses was carried out once at the beginning of the experiment, whereas the EyeLink eye tracker was calibrated prior to each block. Moreover, calibration quality in head-worn systems could also degrade more rapidly over time, at least for older Pupil Labs models and software (Ehinger et al., 2019).

Critically, in addition to finding no significant differences for the SNR effect between the EyeLink and the Pupil Labs eye-trackers, individual participant data correlated between the two devices. This suggests that state-of-the art mobile eye-tracking glasses, such as the Pupil Labs Neon, provide unique opportunities to replace more expensive, stationary eye-trackers with mobile eye-tracking glasses without much loss in sensitivity to assess listening effort. In fact, mobile eye-tracking glasses provide additional benefits by measuring head movements, which also are indicative of listening effort.

### 4.3 Conclusion

The current study compared ocular markers of listening effort between a standard stationary and a modern mobile eye-tracker. Both the stationary and the mobile eye-trackers showed increased pupil sizes and reduced eye movements as listening became effortful, with no significant differences between the devices. Mobile eye□tracking glasses additionally revealed reduced head movements under higher listening demand, which is not available using stationary eye-tracking systems. The current study indicates that mobile eye-tracking reliably captures listening effort, opening a path towards the assessments of listening effort in real life; however, as the present data were collected under constrained laboratory conditions, further validation in more naturalistic environments is warranted.

## Acknowledgments

This work was supported by grants from the Canada Research Chair program (CRC-2023-00383; BH), the Natural Sciences and Engineering Research Council of Canada (RGPIN-2021-02602 to BH; RGPIN-2023-05050 to LKF), and the Canadian Institutes of Health Research (186236; BH).

## Data Availability

The data that support the findings of this study are available from the corresponding author upon reasonable request.

